# Evidence that the Isc Iron-Sulfur Cluster Biogenesis Machinery Delivers Iron for [NiFe]-Cofactor Biosynthesis in *Escherichia coli*

**DOI:** 10.1101/2023.11.17.567542

**Authors:** Alexander Haase, Christian Arlt, Andrea Sinz, R. Gary Sawers

**Affiliations:** Institute for Biology/ Microbiology, Martin-Luther University Halle-Wittenberg, Kurt-Mothes-Str. 3, 06120 Halle (Saale), Germany; Institute of Pharmacy, Center for Structural Mass Spectrometry, Martin-Luther University Halle-Wittenberg, Kurt-Mothes-Str. 3a, 06120 Halle (Saale), Germany

## Abstract

[NiFe]-hydrogenases have a bimetallic NiFe(CN)_2_CO cofactor in their large, catalytic subunit. The 136 Da Fe(CN)_2_CO group of this cofactor is assembled on a distinct HypC-HypD scaffold complex prior to delivery to the apo-catalytic subunit, but the intracellular source of the iron ion is unresolved. Native mass spectrometric (native MS) analysis of HypCD complexes defined the [4Fe-4S] cluster associated with HypD and identified +26 - 28 Da and +136 Da modifications specifically associated with HypC. A HypC_C2A_ variant dissociated from its complex with native HypD lacked all modifications. HypC dissociated from HypCD complexes isolated from *Escherichia coli* strains deleted for the *iscS* or *iscU* genes, encoding core components of the Isc iron-sulfur cluster biogenesis machinery, specifically lacked the +136 Da modification; however, it was retained on HypC isolated from *suf* mutants. The presence or absence of the +136 Da modification on the HypCD complex correlated with the hydrogenase enzyme activity profiles of the respective mutant strains. Notably, the [4Fe-4S] cluster on HypD was identified in all HypCD complexes analyzed. These results suggest that the iron of the Fe(CN)_2_CO group on HypCD derives from the Isc machinery, while either the Isc or the Suf machinery can deliver the [4Fe-4S] cluster to HypD.

## Introduction

[NiFe]-hydrogenases (Hyd) need a bimetallic NiFe(CN)_2_CO cofactor in their catalytic subunit to be able to oxidise or to generate H_2_ [1–3]. Hyd enzymes also have a smaller, electron-transfer subunit that harbours iron-sulphur [Fe-S] clusters, which serve to shuttle electrons to and from the NiFe(CN)_2_CO cofactor. Previous studies carried out by our group showed that the synthesis and insertion of the [Fe-S] clusters into the small subunit of the Hyds in *Escherichia coli* depend on the Isc (iron-<u>s</u>ulphur <u>c</u>luster) biogenesis machinery [4, 5]. The alternative Suf (<u>s</u>ulphur <u>u</u>ptake function) [Fe-S] cluster biogenesis machinery present in *E. coli* is not essential for synthesis of the anaerobically active Hyd enzymes [4].

In contrast to what is understood about [Fe-S] cluster synthesis and insertion into Hyd small subunits, the route of delivery of the iron ion present in the NiFe(CN)_2_CO cofactor is still unclear. The Fe(CN)_2_CO moiety of this cofactor is assembled on a separate scaffold complex [6, 7], requiring the activities of minimally four Hyp (<u>hy</u>drogenase <u>p</u>leiotropy) proteins, HypC, D, E and F [1, 8, 9]. The activities of all four Hyp proteins are necessary for synthesis of the three diatomic ligands (1 x CO and 2 x CN^−^) coordinated to the Fe ion [1–3], but the core of the scaffold complex comprises HypC and HypD. The HypCD complex is proposed to coordinates the Fe(CN)_2_CO group via two conserved cysteine residues: Cys41 on HypD, and the *N*-terminal Cys2 on HypC [10–12]. The HypD protein is redox active and has a conserved and essential [4Fe-4S] cluster, the origin of which is also unclear [10, 11].

Members of the HypC family are small, approximately 10 kDa, proteins that exhibit chaperone-like activity [13, 14]. As well as forming a complex with HypD [11], they also interact with the apo-form of the Hyd catalytic subunit [14, 15]. Using native mass spectrometry (native MS), it was recently shown for a HypC paralogue in *E. coli*, termed HybG, that it interacts specifically with the apo-catalytic subunit of the H_2_-oxidising Hyd-2 enzyme [15]. This corroborates an earlier proposal [14] that the HypC family of proteins (which includes HybG) functions to transfer the Fe(CN)_2_CO group from the HypCD (or HybG-HypD) complex to its cognate apo-catalytic subunit [15, 16]; for example, in *E. coli*, HypC delivers the Fe(CN)_2_CO group specifically to apo-HycE, the precursor of the Hyd forming the H_2_-producing formate hydrogenlyase (FHL) complex [1]. Once this moiety has been introduced into the empty active site cavity, two other Hyp proteins (HypA and HypB) deliver the nickel ion to complete cofactor assembly and insertion [1–3].

It was noted during our native MS analysis of anaerobically isolated *E. coli* HybG-HypD scaffold complexes that HybG carried two post-translational modifications [15]. One of these modifications increased the mass of HybG by +26 Da, while a second modification increased the mass of HybG by a further +136 Da. Both modifications proved to be dependent on the presence of the *N*-terminal Cys residue. The size of this +136 Da modification correlated with the expected size of the Fe(CN)_2_CO group attached to HybG. In further support of this modification representing the Fe(CN)_2_CO group, dissociation of native HybG from a HybG-HypD_C41A_ complex revealed a lack of the +136 Da modification, but retention of the +26 Da modification [15]. *E. coli* mutants synthesising the HypD_C41A_ amino acid variant are incapable of making the [NiFe]-cofactor and thus lack active Hyd enzymes [12]. Together, these findings suggested that the +136 Da modification might represent the Fe(CN)_2_CO group and provided a methodology to identify the origin of the Fe ion in the NiFe(CN)_2_CO cofactor.

Therefore, in the current study, we analysed HypCD complexes isolated from *E. coli* strains with deficiencies in iron transport or [Fe-S] cluster biogenesis. Our findings reveal that the HypC protein dissociated from the anaerobically isolated native HypCD complex carries similar modifications to those identified to be associated with HybG, as well as an additional modification with the mass of a formyl group. Moreover, we show that the appearance of the +136 Da modification is dependent a functional Isc machinery. Finally, our data also show that insertion of the [4Fe-4S] cluster into HypD is not wholly dependent on the Isc machinery.

## Results

### Mutants with defects in iron metabolism synthesise less HypD

Findings of previous studies have shown that *E. coli* mutants with defects in either iron-transport or Isc-dependent [Fe-S] cluster biogenesis lacked, or had severely reduced levels of, Hyd enzyme activity after anaerobic cultivation [4, 5]. Here, we verified and extended these findings, because the main aim of this current study was to examine critically whether the Isc machinery might be a source of the iron for [NiFe]-cofactor biosynthesis. An in-gel hydrogenase activity stain allows simultaneous qualitative visualisation of the activities of Hyd-1, Hyd-2 and Hyd-3 [17]. The results presented in Fig. 1a identified several activity bands attributable to the three Hyd enzymes in an extract derived from the parental strain, MC4100. Furthermore, a H_2_-oxidising activity associated with the formate dehydrogenases (Fdh) N and O could also be identified [see 18]. While an extract derived from a *hypD* deletion mutant lacked activity bands associated with all three Hyd enzymes, the activity associated with Fdh-N and Fdh-O was retained (Fig. 1a). Mutations in *isc* genes encoding components of the scaffold complex (IscU) or the carrier proteins (IscA and ErpA) lacked all H_2_-oxidising enzyme activities (Fig. 1a). In contrast, a mutation in the *sufA* gene, whose gene product usually delivers completed [Fe-S] clusters to specific client protein targets [19], had an activity pattern very similar to the wild-type parental strain. This result is in accord with the Suf machinery not being directly involved in delivery of [Fe-S] clusters to the anaerobic Hyd and Fdh enzymes [4, 5].

**Figure 1.**
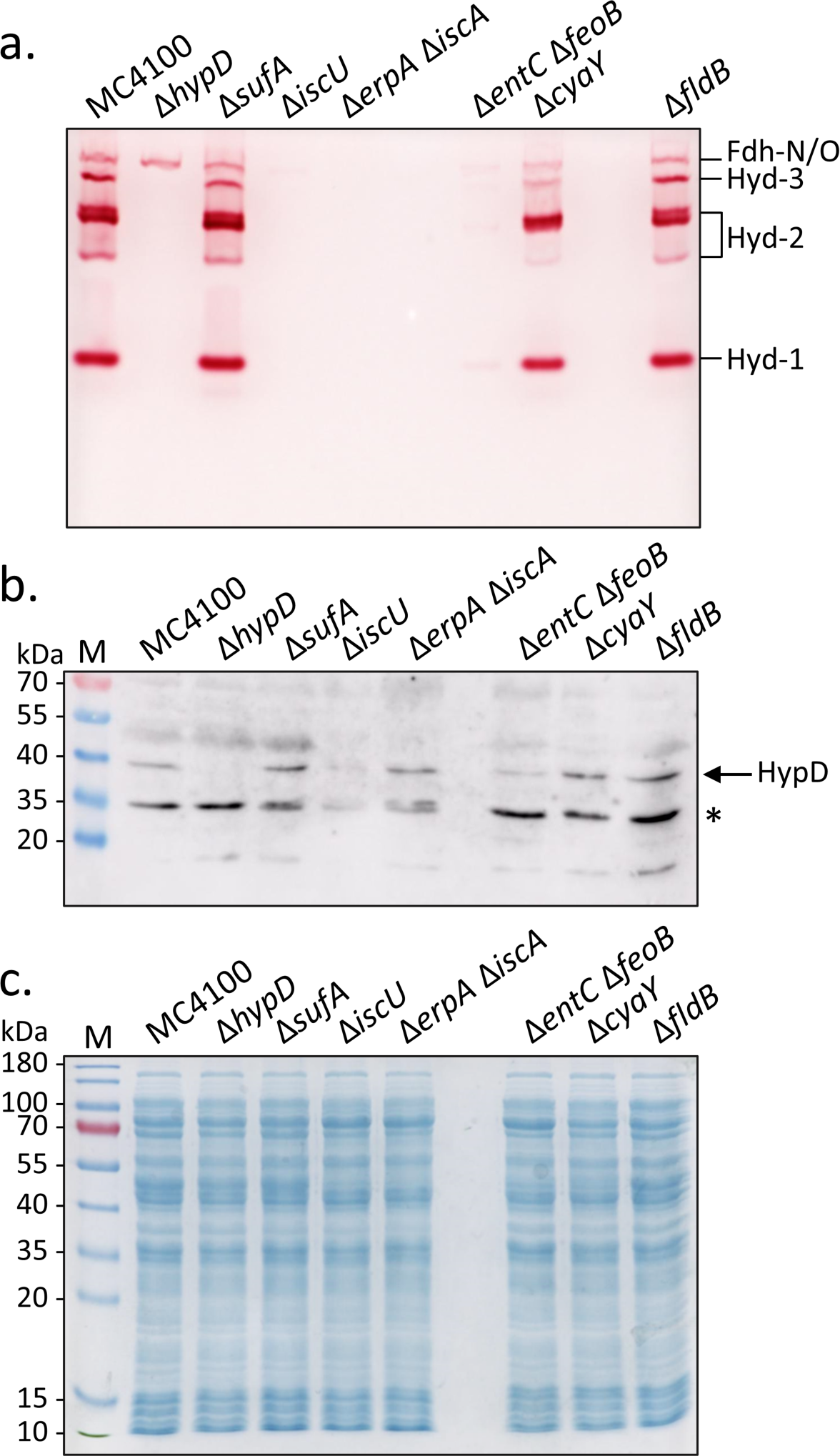
Effects of mutations in key genes of iron metabolism on hydrogenase enzyme activity. a) In-gel hydrogenase enzyme activity determined in extracts of the indicated strains. Aliquots (25 μg of protein) of crude extracts were separated in clear-native polyacrylamide (7.5% w/v) gels. b) Aliquots (25 μg of protein) of the same crude extracts shown in part a) were separated under denaturing conditions in SDS-PAGE (12.5% w/v acrylamide). After transfer to a nitrocellulose membrane, the HypD polypeptide was identified (see arrow) using antiserum raised against HypD (dilution 1:4000). The asterisk denotes an unidentified cross-reacting polypeptide. c) As in part b) but stained with Coomassie-Brilliant-Blue. Molecular mass markers are shown in kDa on the left of panels b and c.

Mutations in genes involved in ferrous (*feo*) and ferric iron (*entC*) uptake have been shown to have a strong negative impact on H_2_oxidising Hyd enzyme activity [20]. Strain CP411, which carries deletions in the genes *feoB* and *entC* had strongly reduced activity of all Hyd enzymes, as well as of the H_2_-oxidising activity associated with Fdh-N and Fdh-O (Fig. 1a). This result indicates that both of these transport routes are likely to be important in providing sufficient iron to the Isc machinery.

Finally, we examined the effect of a deletion mutation in *cyaY,* the bacterial frataxin homologue, and in *fldB*, a potential flavodoxin electron donor for HypCD function. Neither mutation had any significant influence on the H_2_-oxidising enzyme patterns in the respective extracts of their host (Fig. 1a). CyaY is therefore not essential for [Fe-S] cluster biosynthesis [21] or iron delivery during anaerobic hydrogenase enzyme synthesis.

Because HypD has a [4Fe-4S] cluster [11], and due to the fact that the inability to introduce this cofactor into the enzyme results in its destabilisation and degradation [12, 22], it was important to determine whether native HypD levels were affected in the mutants with defective Isc machineries or iron uptake systems. Therefore, the same extracts used to determine Hyd enzyme activities were analysed by western blotting with antiserum raised against HypD (Fig. 1b). Native HypD migrated with a mass of 39 kDa and, although produced at a low level, it could nevertheless be detected in extracts derived from the parental strain MC4100, as well as the Δ*sufA,* Δ*cyaY,* Δ*fldB*, and Δ*erpA-*Δ*iscA* mutants. While still detectable, the level of HypD in the Δ*iscU* and Δ*entC*Δ*feoB* mutants was significantly reduced (Fig. 1b). As expected, an extract derived from the Δ*hypD* mutant, DHP-D, failed to produce a HypD polypeptide and served as a negative control for the experiment. The Coomassie Brilliant Blue-stained gel shown in Fig. 1c confirmed that similar amounts of total protein were applied to each lane. With the possible exception of the Δ*erpA-*Δ*iscA* double-null mutant, CP742, these data revealed a correlation between the absence of Hyd enzyme activity and reduced cellular levels of the [4Fe-4S]-cluster-containing HypD protein.

### Overproduction of the HypCD complex fails to restore Hyd enzyme activity to isc mutants

To determine whether HypD levels limited the manifestation of Hyd activity in the *isc* mutant strains, we introduced multicopy plasmid pT-hypDCStrep and determined the level of HypC- and Hyd-3-dependent FHL activity by measuring H_2_ levels that accumulated in sealed Hungate tubes after overnight growth of each transformed strain (Fig. 2). After anaerobic growth, the parental strain, MC4100, accumulated approximately 30 μmol H_2_ OD_600nm_ ^−1^, while strain DHP-D (Δ*hypD*) failed to produce any H_2_ (Fig. 2), as anticipated [8].

**Figure 2.**
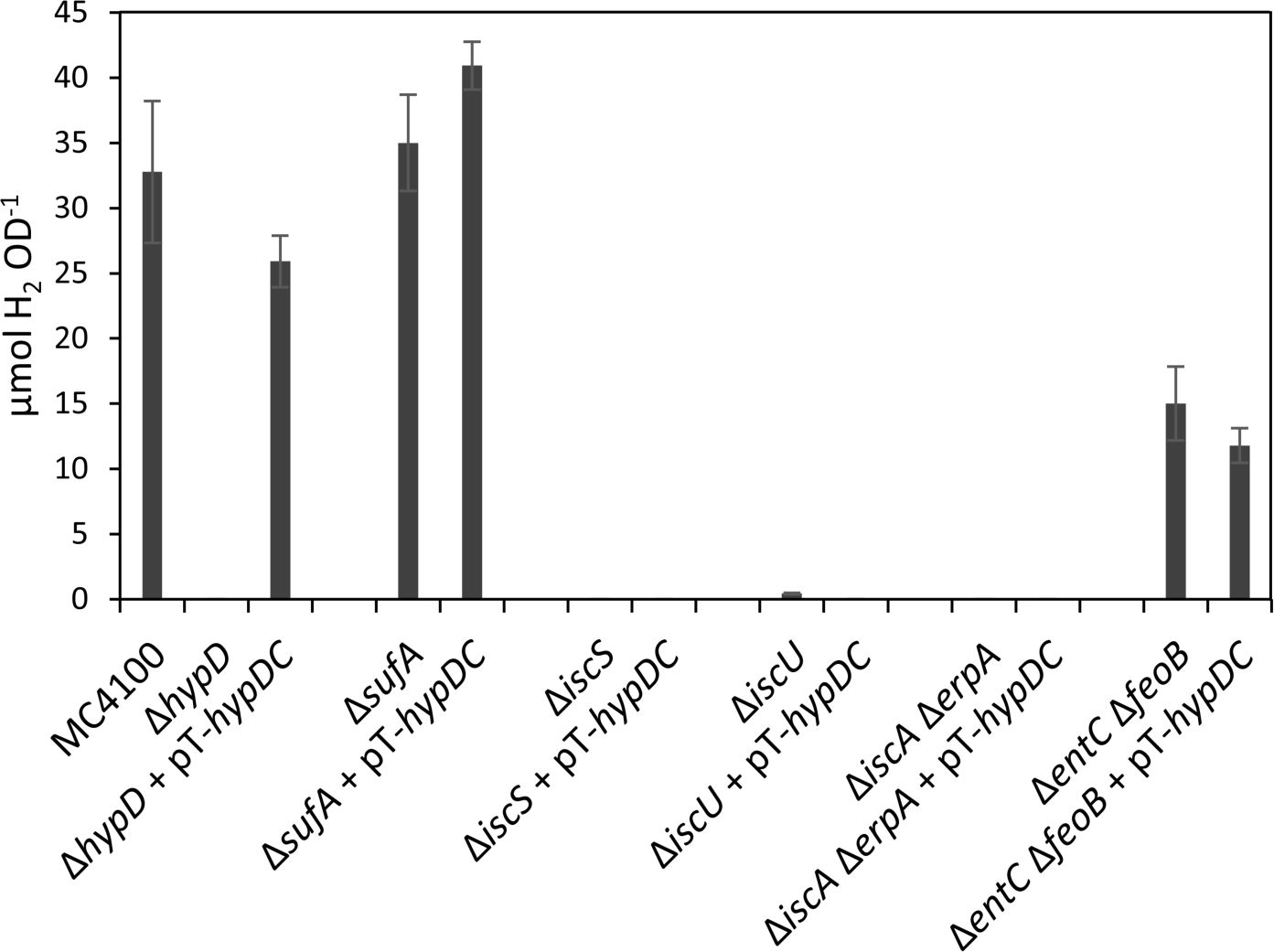
Plasmid-encoded native HypCD complex fails to restore H_2_ evolution to *E. coli isc* mutants. Total hydrogen gas accumulated in the headspace of anaerobic cultures of the indicated strains, with or without plasmid pT-hypDCStrep, and grown for 22h at 37 °C in TGYEP medium is shown.

Introduction of pT-hypDCStrep into DHP-D restored H_2_ production to approximately 80% of the parental level (Fig. 2). While the Δ*sufA* mutant, CP1223, produced H_2_ at a level slightly higher than that of the parental strain, neither the Δ*iscU*, nor the Δ*erpA*Δ*iscA*, nor an Δ*iscS* mutant produced any measurable amount of H_2_ (Fig. 2). Introduction of plasmid pT-hypDCStrep failed to restore H_2_ production to any of these *isc* mutant strains.

The Δ*entC-*Δ*feoB* mutant retained approximately 50% of the H_2_ level accumulated by the parental strain (Fig. 2), which confirms the findings of a previous study [20]. Measurement of the FHL specific activity in exponential-phase, growing cells for the parental strain MC4100 measured an activity of 30 nmol min^−1^ mg protein^−1^, while that for CP411 (Δ*entC-*Δ*feoB*) was 5 nmol H_2_ min^−1^ mg protein^−1^. This low-level FHL activity indicates that a few active FHL complexes were synthesised in the mutant.

### HypD isolated in complex with HypC from isc mutants retains a [4Fe-4S] cluster

StrepII-tagged HypCD complexes were isolated under anoxic conditions from various mutants that had been transformed with the plasmid pT-hypDCStrep; henceforth these complexes will be referred to as HypCD throughout this study and it is important to note that HypC carried a StrepII tag in all of the experiments shown. All anaerobically isolated complexes from all strains that were analysed had a similar yellow-brown colour (data not shown, but see below), which is characteristic of a FeS protein. Analysis of the isolated HypCD complexes by SDS-PAGE followed by Coomassie Brilliant Blue-staining revealed that they all showed minimally 90% purity (Fig. 3a). The HypD polypeptide in complexes isolated from the *iscU* single mutant and the *erpA-iscA* and *entC-feoB* double-null mutants had an additional, weak polypeptide that migrated slightly faster (∼39 kDa) than the main HypD polypeptide. This additional polypeptide has been observed previously, but with greater intensity, in HypD amino acid variants lacking a complete [4Fe-4S] cluster [22]. Western blot analyses of the same samples confirmed that the dominant polypeptide migrating at ∼39 kDa was HypD (Fig. 3b) and the polypeptide migrating at ∼10 kDa was StrepII-tagged HypC (Fig. 3c).

**Figure 3.**
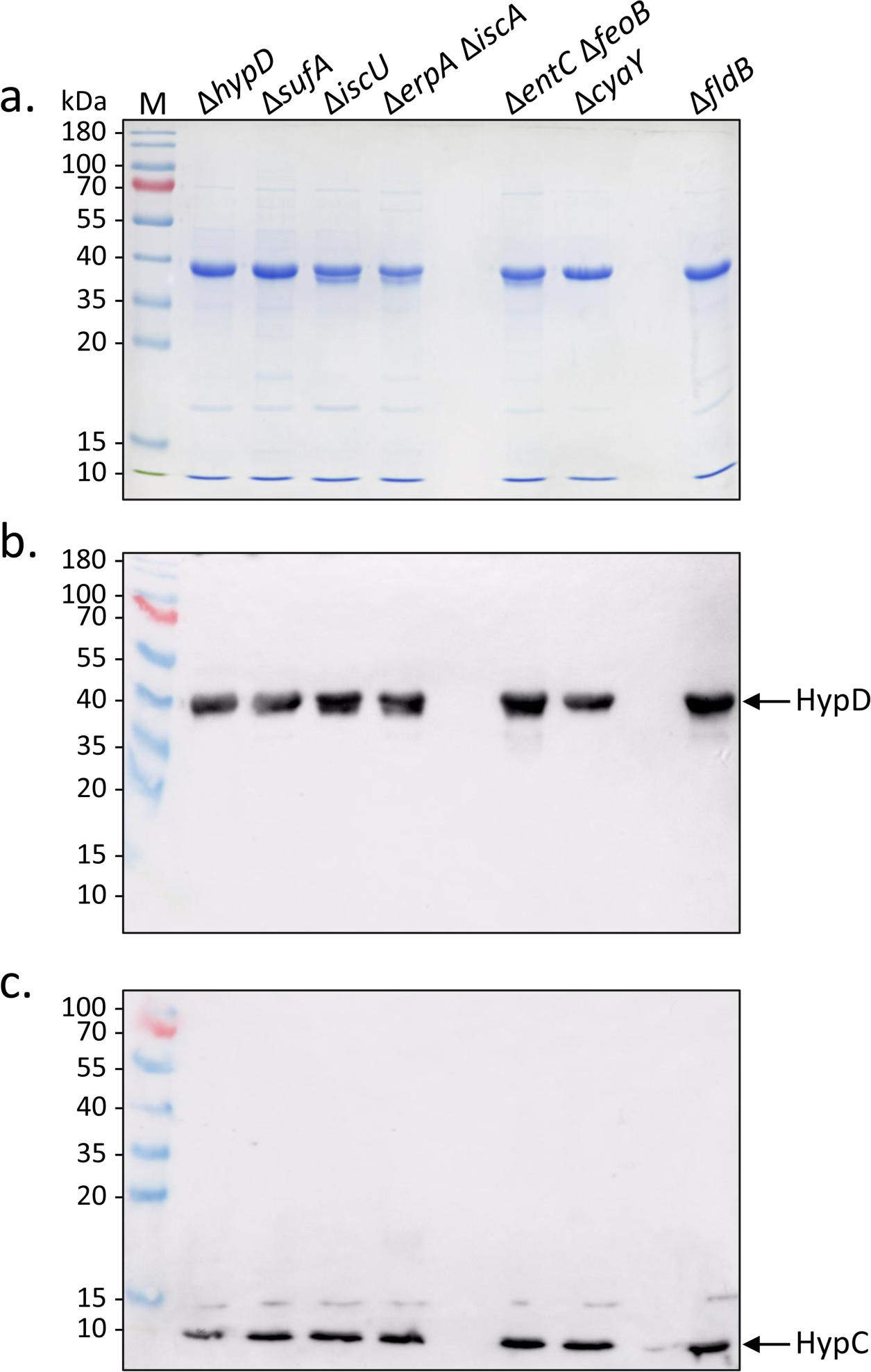
Purified StrepII-tagged HypCD complexes isolated from strains with defects in iron metabolism. a) Coomassie-Brilliant-Blue-stained SDS-PAGE (12.5% w/v polyacrylamide) of purified StrepII-tagged HypCD complexes (5 μg protein) isolated from the indicated strains. b) A gel identical to that shown in part a), but challenged with antiserum containing antibodies specific for HypD (diluted 1:4000). c) A gel similar to that shown in part a), but with 7.5 μg of protein applied per lane and challenged with antiserum containing antibodies against HypC (diluted 1:4000). Molecular mass markers in kDa are shown on the left of each panel. Arrows identify the migration positions of HypD or HypC.

Analysis of these anaerobically isolated complexes by UV-vis spectroscopy revealed a absorption broad shoulder between 380 nm and 420 nm for the native HypCD complex isolated from strain DHP-D carrying plasmid pT-hypDCStrep (Fig. 4a). This absorption shoulder is a characteristic feature of HypD within these complexes [11, 22]. The spectrum of the HypCD complex isolated from the Δ*sufA* mutant, CP1223 carrying plasmid pT-hypDCStrep (Fig. 4b), was almost identical to that of the complex isolated from DHP-D/ pT-hypDCStrep, while the broad shoulder in the spectra of the native complexes isolated from the *iscU* (Fig. 4a) and the *erpA-iscA* (Fig. 4b) mutants was weaker in intensity. Spectra of the native HypCD complex isolated from the Δ*cyaY* and Δ*fldB* mutants transformed with plasmid pT-hypDCStrep had near-identical features to the complex isolated from the Δ*hypD* mutant, DHP-D carrying plasmid pT-hypDCStrep, while the shoulder in the spectrum of the complex isolated from the iron transport-deficient mutant, CP411 carrying plasmid pT-hypDCStrep, was only marginally weaker (data not shown). Together, these data indicate that all anaerobically isolated HypCD complexes retained the spectroscopic features of a [4Fe-4S] cluster.

**Figure 4.**
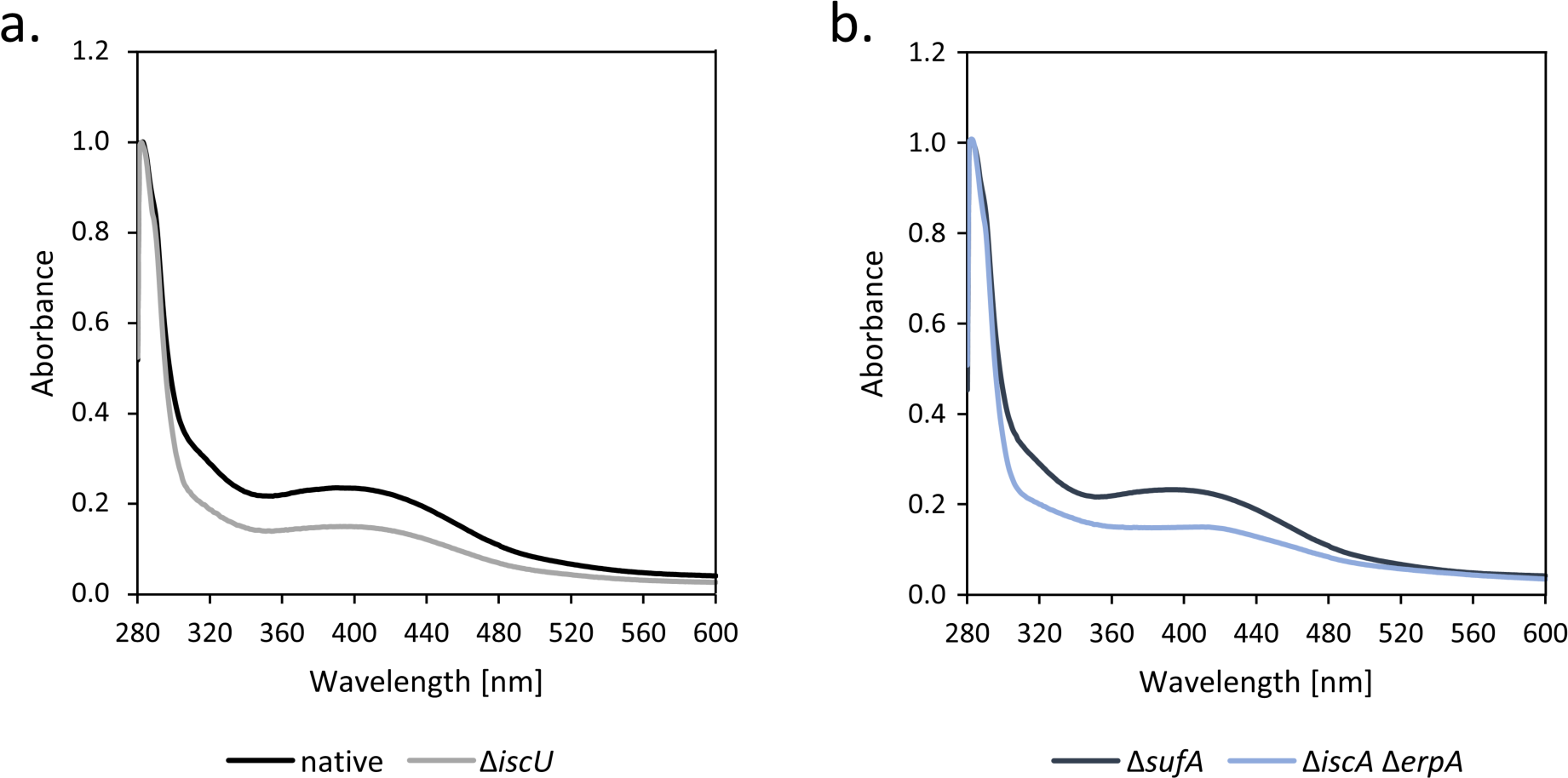
UV-vis spectroscopy indicates HypD contains a [4Fe-4S] cluster. Shown are UV-vis spectra of StrepII-tagged HypCD complexes (1 mg ml^−1^) isolated from the strains indicated. a) HypCD isolated from DHP-D (Δ*hypD*) transformed with pT-hypDCStrep (blue spectrum) or from CP1244 (Δ*iscU*) transformed with pT-hypDCStrep (red spectrum); b) HypCD isolated from CP1233 (Δ*sufA*) (black spectrum) transformed with pT-hypDCStrep, or CP742 (Δ*iscA-*Δ*erpA*) transformed with pT-hypDCStrep (green spectrum).

### Modifications associated with the HypC and HypD proteins determined by native MS

Native MS revealed that the majority of the anaerobically isolated HypCD complexes primarily had a 1:1 stoichiometry (Fig. 5a, upper panel; charge-state distribution +9 to +14), but minor amounts of HypCD complexes with stoichiometries of 2:1 and 2:2 were also identified (Figure 5a upper panel; peaks labelled * and #). Signals attributable to minor amounts of the free, monomeric species of StrepII-tagged HypC (+4 and +5 charged species) and HypD (+10, +12 and +13 charged species) could be identified. It is likely that these dissociated species are also present in solution. The isolated HypCD complexes showed nearly identical behavior with respect to stoichiometry and charge state distribution in native MS, regardless of the genotype of the *E. coli* strain in which they were synthesised (Fig. 5b). Consequently, the interaction between HypC and HypD was independent of the individual extent of any modifications on either protein. This conclusion was substantiated by the isolation and analysis of a HypD-HypC(C2A) complex in which HypC had an amino acid exchange of C2A (Fig. 5a, lower panel). The spectrum revealed a similar pattern of charge state distribution and complex stoichiometry with only some variation in the respective intensities. This result demonstrates that the *N-*terminal cysteine residue of HypC is not required for its interaction with HypD.

**Figure 5.**
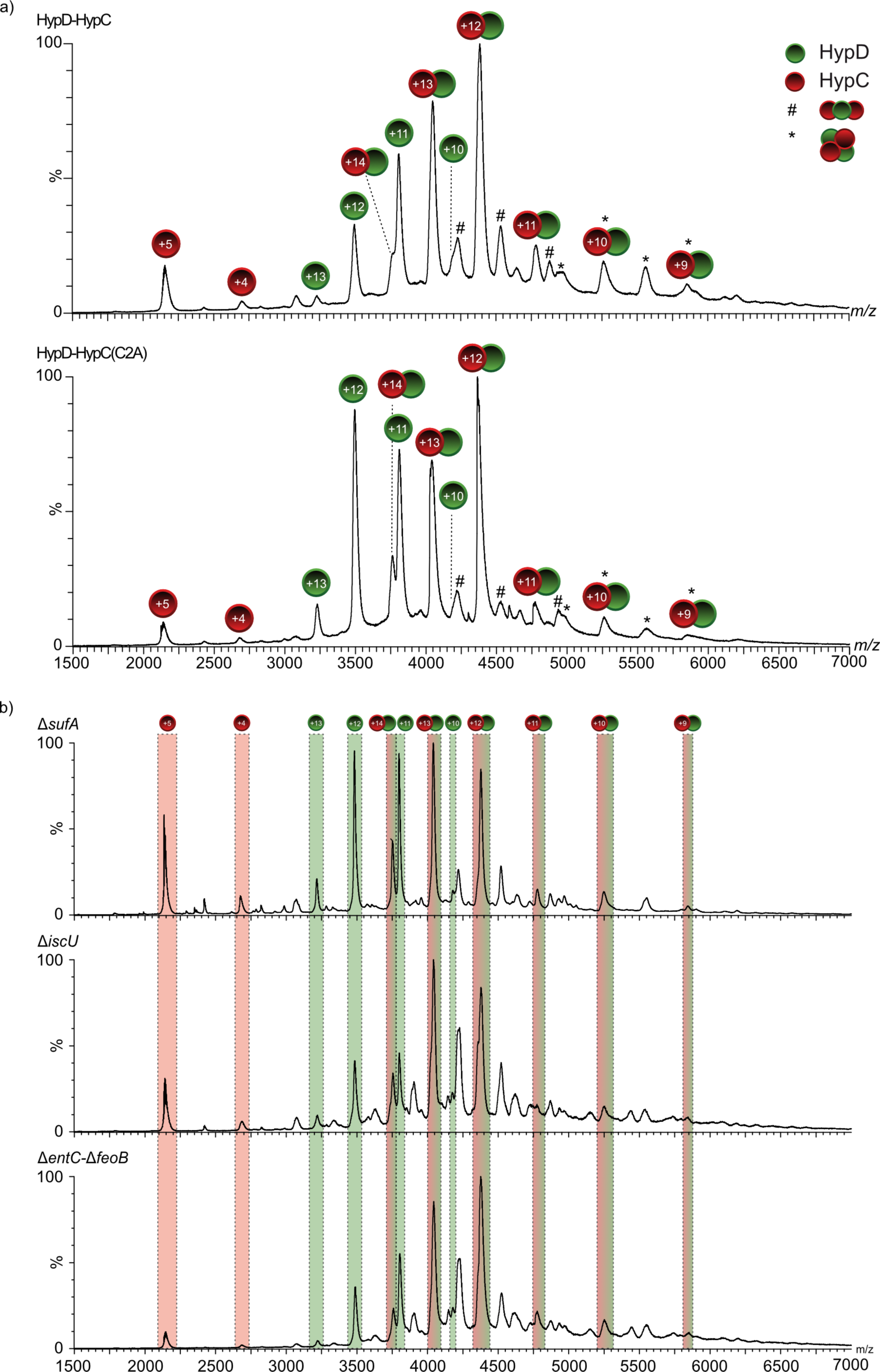
Native mass spectra of HypCD complexes isolated from different *E. coli* mutants. a) Native mass spectrum of native HypCD complex (upper panel) and HypC_C2A_D complex (lower panel) isolated from strain DHP-D (Δ*hypD*) transformed with pT-hypDCStrep or pT-hypDC(C2A)Strep, respectively, and analysed at a collision energy of 30V. Stoichiometry of the main complex species is StrepII-HypC:HypD of 1:1 (indicated by red circles for HypC and green circles for HypD), with minor species showing 2:1 (#) and 2:2 (*) ratios. b) Native mass spectra of the HypCD complex isolated from strains CP1233 (Δ*sufA*), CP1244 (Δ*iscU*) or CP411 (Δ*entC-*Δ*feoB*). All strains carried plasmid pT-hypDCStrep. Signals corresponding to HypC (red overlay), HypD (green overlay) and the 1:1 complex (red-green overlay) are labeled accordingly.

Dissociating the HypCD complex using tandem MS/MS experiments (collision-induced dissociation, CID-MS/MS) of the isolated +12 charged species yielded signals for charged HypC species (red circles, charge states +4 through +6, Figure 6a) and signals corresponding to HypD (green circles, charge states +6 though +8), with the +7 charge-state giving the strongest signal (Figure 6a). Enlarging the *m/z* region of the dissociation products with the highest intensity for HypC (Fig. 6b; *m/z* region = 2115-2170; +5 charged species) and HypD (Fig. 6c; *m/z* region = 5900-6000; +7 charged species) revealed several adduct signals for both proteins. The species of HypD with a *m/z* value of 5960.9, corresponding to a mass of 41715 Da, indicated that this species included the [4Fe-4S] cluster (41363.4 Da + 351.6 Da; Fig. 6c). The adjacent signals indicated successive loss of sulphur, with the signal of *m/z* value 5941.8 corresponding to HypD with only [4Fe] of the [4Fe-4S] cluster remaining. Notably, all isolated HypCD species, regardless from which mutant they were prepared, showed the same HypD signal distribution as the wild type (Fig. 6c; Fig. S1) and retained near-native ATPase enzyme activity (Fig. S2), which supports our contention that the HypD component of these complexes had a complete, redox-active [4Fe-4S] cluster [22].

**Figure 6.**
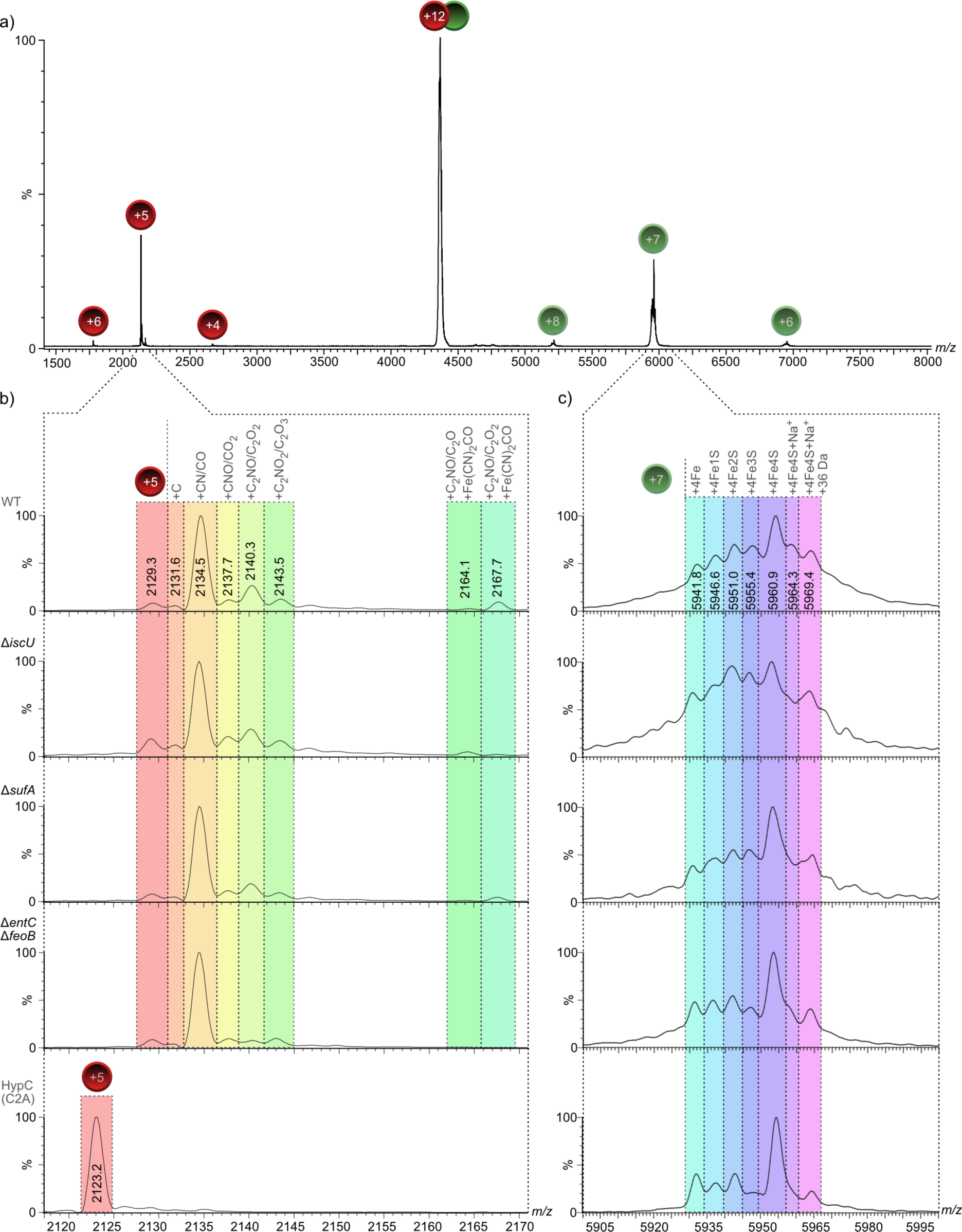
Native MS spectra of HypC and HypD dissociated from HypCD complexes reveal absence of the +136 Da modification on HypC in *isc* mutants but presence of the [4Fe-4S] cluster on HypD. a) Mass spectrum of the dissociation of the +12 charged ion species of the HypCD heterodimer into HypC (charge states +4 through +6, red spheres) and HypD (charge states +6 through +8, green spheres) at collision energy of 90V isolated from strain DHP-D transformed with plasmid pT-hypDCStrep as positive control. b) Zoom in for HypC (charge state +5), as well as c) zoom in for HypD (charge state +7) are shown for isolated complexes from strains transformed with pT-hypDCStrep: DHP-D (“WT”), CP1233 (Δ*sufA*), CP1244 (Δ*iscU*) and CP411 (Δ*entC-feoB*), and HypC_C2A_ dissociated from the HypC_C2A_HypD complex isolated from strain DHP-D (Δ*hypD*) transformed with pT-hypDC(C2A)Strep. Signals are labeled with the corresponding *m/z* value in the first zoomed row. Potential modifications are indicated above the colored overlay. Note that a putative methyl thiazolidine modification accounting for the +26 Da species is not indicated in the Figure.

The dissociation of HypC from the HypCD complex revealed additional signals for the +5 charge species starting at a *m/z* value of 2129.3 (Fig. 6b). Like family members from other archaeal and bacterial species, HypC is known to lose the formyl-methionine after translation [1], indicating that the 2129.3 species represents native, otherwise unmodified, HypC lacking M1 and carrying a *C*-terminal StrepII tag (+5 charge state, deconvoluted mass: 10,641.5 Da). This was confirmed by CID-MS/MS analysis of the HypC_C2A_-HypD complex (Fig. 6b, bottom spectrum), which revealed a single, major species with a *m/z* value of 2123.2 (+5 charge state, deconvoluted mass: 10,611 Da). Notably, the other modified species of HypC were lost by the C2A conversion (Fig. 6b), suggesting that all modifications were associated with, or dependent on, the *N*-terminal cysteine residue. HypC dissociated from HypCD complexes isolated from all other genetic backgrounds (Fig. 6b, WT, Δ*iscU,* Δ*sufA,* Δ*entC* Δ*feoB)* appears to be present in the same modification states, with the highest abundance species having a *m/z* value of 2134.5 (+5 charge state, deconvoluted mass: 10,667.5 Da) with an additional mass of approximately 26 Da. (Fig. 6b). A similar modification was identified recently for StepII-tagged HybG [15].

The species at *m/z* 2140.3 corresponded to StrepII-tagged HypC with an additional mass of 55 Da, which suggests the addition of a CO (or formyl) molecule (28 Da), together with the +26 Da modification. The less abundant species with *m/z* values of 2137.7 and 2143.5 each indicated an additional mass of 16 Da when compared with their respective preceding signals (2134.5 and 2140.3), suggesting oxidation events on the protein (Fig. 6b). Finally, the species with a *m/z* value of 2167.9 corresponded to HypC carrying the +26 Da modification, the +28 Da modification, along with a further +136 Da modification (Fig. 6b, cyan overlay at right). A similar modification (+136 Da) was identified previously on StrepII-tagged HybG that proved to be dependent on the essential Cys41 in HypD and was proposed to represent the Fe(CN_2_)CO species [15]. An overview of the potential identities of the adducts on HypC is shown in Fig. S3.

### The “+136 Da” modification on HypC is absent in mutants with a defective core Isc machinery

To determine whether the observed modifications on HypC depend on the Isc or Suf iron-sulphur cluster biogenesis machineries of *E. coli*, we isolated the StrepII-tagged native HypCD complex from a variety of mutants with defects in cellular iron metabolism but carrying plasmid pT-hypDCStrep (Fig. 6b; Fig. S4). The +26 Da modification on the HypCD complex could be identified in complexes isolated from all of the mutants used is this study. In contrast, dissociation of HypC from complexes isolated from *iscU, iscS* and *entC-feoB* mutants either lacked, or had a strongly reduced intensity of, the peak corresponding to the “+136 Da” modification, while complexes isolated from *cyaY*, *fldB*, *sufA* and *iscA-erpA* mutants all retained the modification on HypC (Fig. 6b and Fig. S4). These data reveal a broad correlation between the presence of the “+136 Da” modification, which we consider to represent the Fe(CN)_2_CO group, and the synthesis of active hydrogenases (see Fig. 1a). The exceptions were HypC dissociated from the complex that had been isolated from the *entC-feoB* mutant, which retained some ability to accumulate H_2_ gas, but HypC showed no or very low levels of the “+136 Da” modification, and HypC isolated from the *iscA-erpA* double null mutant, which retained the +136 Da modification, but the mutant had no Hyd activity (see Fig. 1a and Fig. 2). As indicated above, low amounts of FHL complexes can account for retention of low-level H_2_ production in iron-transport mutants [20], and an inability of *iscA-erpA* mutants to deliver FeS cluster to the electron-transferring subunits of Hyd enzymes obviates activity despite maturation of the catalytic subunit [5].

## Discussion

The identification of a similar set of chemical modifications on HypC to those recently identified on its paralogue, HybG [15], is strong evidence that these modifications represent either intermediates during biosynthesis of the Fe(CN)_2_CO moiety, or bound metabolic precursors of the diatomic ligands. The current study demonstrates unequivocally that the *N*-terminal cysteine residue is essential for manifestation of all modifications identified to be associated with native HypC. The +26 Da modification (*m/z* 2134.8) is the most abundant of all the modifications and is independent of HypD, because it is also present on HypC and HybG isolated from a Δ*hypBCDE* mutant [reported in 15]. In contrast, the modifications with *m/z* values of 2140.5, 2143.7 and 2167.9 (sequential +28.5, +44.5, and +136 Da modifications added to the +26 Da modified species) likely result from either the further oxidation of a formyl group (+ 28.5 Da; *m/z* 2140.5) to deliver the +44.5 Da species, or from the direct binding of CO_2_ (*m/z* 2143.5) [15, 23]; the +136 Da deconvoluted mass increase is in accord with the Fe(CN)_2_CO species [15].

The mass of 26 Da could be accounted for by a thiocyanate (see Fig. 6b), or by a methyl thiazolidine cyclisation; however, with a mass accuracy in this measurement range of +/− 0.7 Da, this makes a *N*-terminal CO modification unlikely (see Fig. 6b and Fig. S3). Notably, as the +26 Da modification is also observed in a strain lacking a HypE enzyme (C. Arlt, R. G. Sawers, unpublished observation), this rules out that any potential thiocyanate modification is derived from HypEF [24]. Although not depicted in the summary of potential adducts presented Fig. S3, a methyl thiazolidine modification (+26 Da) could also potentially result from interaction of cysteine with metabolically-derived acetaldehyde, or by derivatisation of a *S*- or *N*-acetyl group, which might suggest that it arose during preparation or analysis [25, 26]. The carboxylate group of a thio- or *N-*acetyl group bound to the *N*-terminal cysteine residue of HypC has been suggested previously as a possible intermediate in CO synthesis [27].

While the +26 Da and +136 Da modifications are found on both HypC and HybG, HypC has an additional +28 Da modification not previously identified on HybG [15]. This modification would accord with a formyl adduct, which has been shown to be an intermediate during HypX-dependent aerobic synthesis of CO in *Ralstonia eutropha* [28]. Despite all three modifications being dependent on the *N*-terminal cysteine residue, the question arises as to where the +136 Da species is located on HypC. Notably, the +136 Da modification has not been observed in the absence of the +26 Da modification; however, as shown in this study and previously [15], the +26 Da species is independent of the +136 Da modification. Future labelling studies will be required to define both the precise chemical nature and the location on HypC of this species.

The new and somewhat unexpected finding of the current study was the demonstration that, despite HypD retaining a [4Fe-4S] cluster in all HypCD complexes isolated from different mutants with defects in cellular iron metabolism, the +136 Da species was absent when the HypCD complex was isolated from either *iscU* or *iscS* mutants, while the other modifications were retained on HypC. These data indicate that the [4Fe-4S] cluster on HypD can presumably be introduced by the Suf machinery when the Isc system is inactivated. The fact that the [4Fe-4S] cluster was present on HypD was demonstrated by UV-vis spectroscopy, by native-MS and was supported by the fact that these isolated complexes retained a native ATPase activity, which is not the case when HypD lacks its [4Fe-4S] cluster [22].

Furthermore, these data suggest that, in contrast to a previous proposal [11], the Fe ion that ultimately binds the diatomic ligands is unlikely to be cannibalised from the [4Fe-4S] cluster on HypD, because the cluster is retained regardless of whether Suf or Isc is defective, while the +136 modification is lost in the absence of the core Isc proteins. This argument is further supported by the fact as the +136 Da modification was still partially retained when HypCD was isolated from the mutant lacking the [Fe-S] cluster-carrier and delivery proteins IscA and ErpA, which suggests that the iron might derive directly from either IscS or more likely IscU [19].

Recent phylogenomic analyses [29] have revealed that iron-sulphur cluster biosynthesis machineries are of ancient origin, which is in line with their involvement in delivery of the iron for [NiFe]-cofactor biosynthesis. Moreover, as well as the identification of two new minimal [Fe-S] cluster assembly machineries, one of which is related to Suf and is suggested to have been present in the ‘last universal common ancestor’, that study has also revealed that the Suf machinery is considerably more widespread than Isc in archaeal and bacterial genomes [29]. As many microorganisms have only the Suf machinery, this suggests that this machinery might also be used for iron delivery to the Hyp machinery in those microorganisms. Future studies will be required to determine whether this is indeed the case and to identify which components of the Isc or Suf machineries are responsible for delivery of the iron ion to Hyp.

## Materials and Methods

### Bacterial strains, plasmids and growth conditions

The *E. coli* strains used in this study are listed in Table 1. MC4100 (F^−^, *araD139,* Δ(*argF-lac*)*U169,* λ^−^, *rpsL150, relA1*, *deoC1, flhD5301*, Δ(*fruK-yeiR*)*725*(*fruA25*), *rbsR22*, Δ(*fimB-fimE*) [30], its isogenic mutant derivatives, DHP-D (Δ*hypD*) [8]. The plasmids used included pT-hypDCStrep [11], and pT-hypDEFC_C2A_Strep, which was created by substituting the codon encoding cysteine at amino acid residue position 2 on HypC with a codon decoding as alanine using site-directed mutagenesis (Q5 Site-Directed Mutagenesis Kit, New England Biolabs) employing the oligonucleotides HypC_fwd_ (5’-TATACATATG<u>GCG</u>ATAGGCGTTCCCGG-3’) and HypC_rev_ (5’-TCTCCTTCTTAAAGTTAAACAAAATTATTTC-3’). *E. coli* strain XL1-Blue was used for standard cloning procedures [32].

**Table 1.**
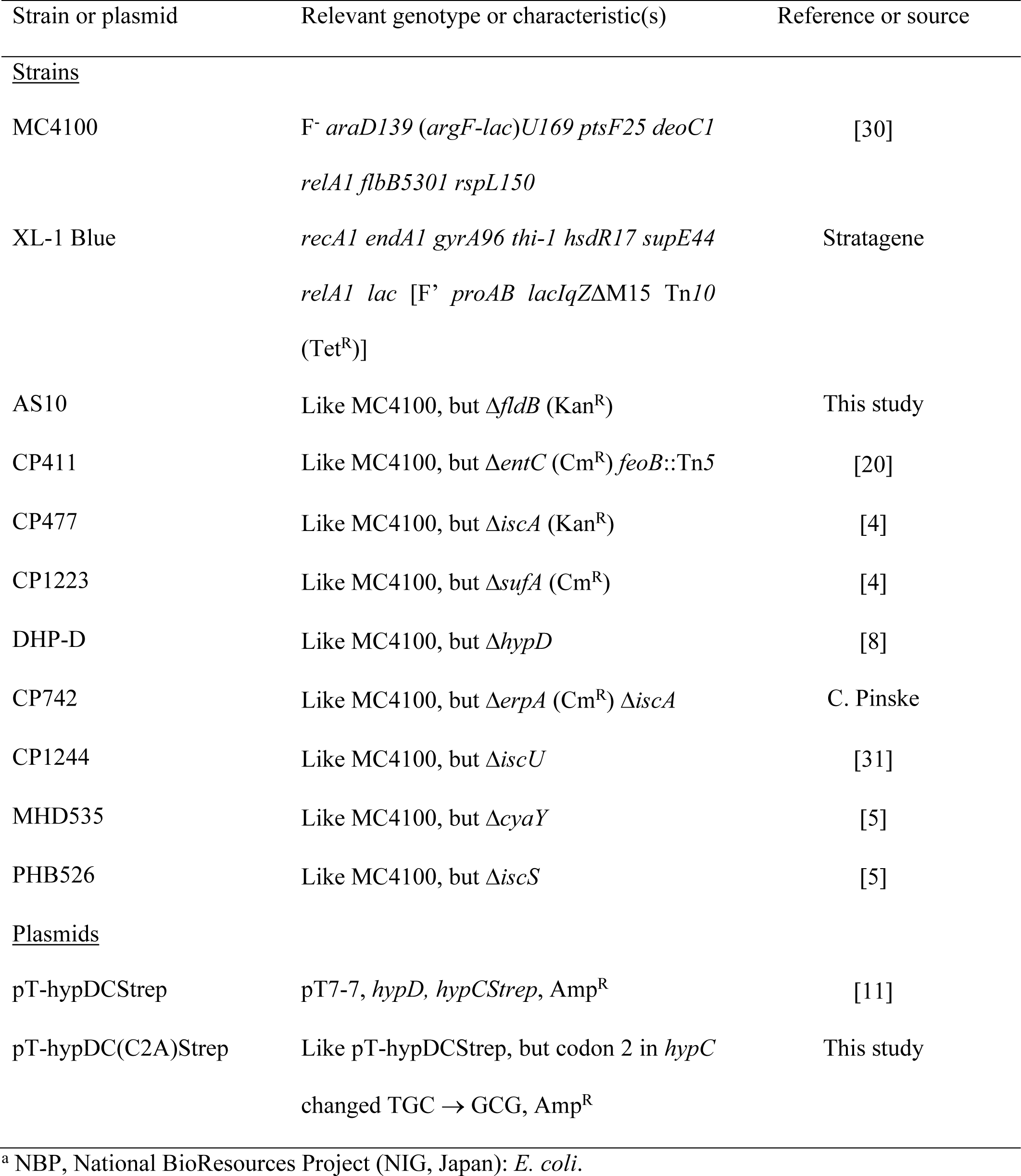
Strains and plasmids used in this study.

| Strain or plasmid | Relevant genotype or characteristic(s) | Reference or source |
| --- | --- | --- |
| <u>Strains</u> |  |  |
| MC4100 | *F<sup>-</sup> araD139 (argF-lac)U169 ptsF25 deoC1*<br>*relA1 flbB5301 rspL150* | [30] |
| XL-1 Blue | *recA1 endA1 gyrA96 thi-1 hsdR17 supE44*<br>*relA1 lac [F' proAB lacIqZΔM15 Tn10*<br>*(Tet<sup>R</sup>)]* | Stratagene |
| AS10 | Like MC4100, but Δ*fldB* (Kan<sup>R</sup>) | This study |
| CP411 | Like MC4100, but Δ*entC* (Cm<sup>R</sup>) *feoB::Tn5* | [20] |
| CP477 | Like MC4100, but Δ*iscA* (Kan<sup>R</sup>) | [4] |
| CP1223 | Like MC4100, but Δ*sufA* (Cm<sup>R</sup>) | [4] |
| DHP-D | Like MC4100, but Δ*hypD* | [8] |
| CP742 | Like MC4100, but Δ*erpA* (Cm<sup>R</sup>) Δ*iscA* | C. Pinske |
| CP1244 | Like MC4100, but Δ*iscU* | [31] |
| MHD535 | Like MC4100, but Δ*cyaY* | [5] |
| PHB526 | Like MC4100, but Δ*iscS* | [5] |
| <u>Plasmids</u> |  |  |
| pT-hypDCStrep | pT7-7, *hypD*, *hypCStrep*, Amp<sup>R</sup> | [11] |
| pT-hypDC(C2A)Strep | Like pT-hypDCStrep, but codon 2 in *hypC*<br>changed TGC → GCG, Amp<sup>R</sup> | This study |

<sup>a</sup> NBP, National BioResources Project (NIG, Japan): *E. coli*.

*E. coli* strain DHP-D (Δ*hypD*) cultivated in modified TB medium (2.4% w/v yeast extract, 1.2% w/v peptone from casein, 0.04% w/v glycerol, 0.4% w/v glucose and 0.003% w/v magnesium sulfate heptahydrate) [23] was used to overproduce native StrepII-tagged HypC-HypD and StrepII-HypC_C2A_-HypD protein complexes for analysis of HypC modifications. To test the effects of mutations in genes encoding proteins required for iron metabolism on the HypC modifications and HypD [4Fe-4S] cluster content, the respective strains (see Table 1) were transformed with plasmid pT-hypDCStrep using standard procedures [32].

Growth of strains for determination of hydrogenase enzyme activity and in-gel enzyme activity staining after native PAGE was performed at 37 °C in standing liquid cultures in the buffered rich medium TGYEP (1% w/v tryptone, 0.5% w/v yeast extract, 0.8% w/v glucose, 100 mM potassium phosphate, pH 6.5) [22].

Cultivation of strains for protein purification was done at 30 °C anaerobically in modified TB medium until an optical density at 600 nm of between 1.0 and 1.2 was attained. The growth medium included 100 μg ml^−1^ of ampicillin to maintain plasmid selection. Cells were harvested by centrifugation of the culture for 15 min at 50,000 g at 4 °C and washed cell pellets were either used immediately or stored at −20 °C until use.

### Protein purification

All steps in cell disruption and protein purification were carried out under anoxic conditions in an anaerobic chamber (Coy Laboratories, Grass Lake, USA). Purification of StrepII-tagged HypC-HypD complexes was carried out exactly as previously documented for StrepII-tagged HybG-HypD complexes using StrepTactin Sepharose^®^ (IBA, Göttingen) [15, 23]. When required, eluted, pooled protein fractions from the affinity chromatography steps were immediately buffer-exchanged into anaerobic 50 mM Tris-HCl, pH 8, using 5 ml PD-10 columns containing G-25 Sephadex matrix (Cytiva) and then chromatographed on a 1 ml Q-Sepharose^®^ fast-flow column (Cytiva) equilibrated with the same buffer. Bound proteins were eluted stepwise using equilibration buffer containing 50, 150, 300 and 500 mM NaCl [22].

After buffer-exchange into anaerobic 50 mM Tris-HCl, pH 8, containing 150 mM NaCl (buffer A), protein samples were concentrated using Amicon centrifugal concentration filters (cut-off or 50 kDa) and samples were stored at −80°C.

Protein concentration was determined as described [33].

### Measurement of H_2_ production

The amount of H_2_ gas that accumulated in the gas phase of Hungate tubes after anaerobic growth of strains in TGYEP medium was performed exactly as described [31].

### Determination of ATPase activity of HypCD complexes

The ATPase enzyme activity associated with HypCD complexes was determined exactly as described [22, 34].

### Nondenaturing PAGE and hydrogenase activity staining

Non-denaturing PAGE (polyacrylamide gel electrophoresis) was performed according to [17]. Aliquots (25–50 μg of protein) of crude extracts were separated using gels that included 7.5% (w/v) polyacrylamide and 0.1% (w/v) Triton X-100. Before gel application, the crude extracts were incubated with a final concentration of 4% (v/v) Triton X-100 at 4 °C for 15 min. Visualization of H_2_-oxidizing activity of Hyd-1, Hyd-2, and Hyd-3 was performed as described previously [17], whereby gels were incubated overnight at 25 °C in an atmosphere of 95% N_2_ : 5% H_2_. Experiments were repeated minimally three times using biological replicates, and a representative gel is shown.

### Denaturing polyacrylamide gel electrophoresis and western blotting

Polypeptides in cellular extracts or in purified HypCD complexes were analysed by sodium dodecylsulphate polyacrylamide gel electrophoresis (SDS-PAGE), as described [35]. In the current study, gels containing 12.5 % (w/v) polyacrylamide were used. After electrophoretic separation, polypeptides were visualised by staining with Coomassie Brilliant Blue G250 (Sigma-Aldrich, Germany), or western blotting. Transfer to nitrocellulose membranes, treatment with antiserum raised against either HypC or HypD and subsequent visualization of signals was done as previously described [22].

### UV–vis spectroscopy

The spectral properties of the purified StrepII-tagged HypC-HypD complexes were analyzed in the wavelength range 280–600 nm using a Shimadzu UV-1900i UV–vis spectrophotometer (Shimadzu Europe GmbH, Duisburg, Germany) and quartz cuvettes with a 1 cm pathlength. The protein concentration used to record the spectra was typically 1 mg mL^−1^ [22].

### Mass spectrometry analyses

For native MS measurements the buffer was exchanged to 500 mM ammonium acetate, pH 6.8, by rapid online buffer exchange as described previously [15].

The concentration of protein solutions after buffer-exchange was approximately 10 μM. Native MS was carried out on a High-Mass Q-TOF II instrument (Waters Micromass / MS Vision) equipped with a nano-electrospray ionization (ESI) source. The applied capillary voltage ranged from 2.0 to 2.3 kV, while the sample cone voltage varied from 100 to 160 V. The source pressure was adjusted to 10 mbar and the pressure in the collision cell was adjusted to 10^−2^ to 2*10^−2^ mbar. MS measurements were carried out using MS profile mode for the quadrupole to guide ions within the *m/z* region of interest. The acceleration voltage in the collision cell was set to 30V for MS measurements. Dissociation experiments were carried out by collision-induced dissociation (CID) for the selected ion species. To achieve dissociation of protein complexes the collision energy was set to 90V. Data were recalibrated by using cesium iodide (CsI).

## Acknowledgments

We thank Dr. Constanze Pinske for providing strains and for valuable discussions. This work was supported by the Deutsche Forschungsgemeinschaft (DFG) through the SPP1927 “Iron-sulfur for life” initiative to RGS and the RTG 2467, project number 391498659 “Intrinsically Disordered Proteins—Molecular Principles, Cellular Functions, and Diseases” to AS.

## Author contributions

AH, CA, AS and RGS designed the experiments and analysed the data. AH and CA carried out all the experiments. AS and RGS drafted the manuscript and conceived the study. All authors read and approved the final version of the manuscript.

## Data Availability Statement

All data are either presented in this manuscript or are freely available from the corresponding author upon request.

## Additional Information

All authors declare that they have no conflict of interest.

## Notes

### Competing Interest Statement

The authors have declared no competing interest.

