## Supplementary Figures 1-3 for "Evidence that the Isc Iron-Sulfur Cluster Biogenesis Machinery Delivers Iron for [NiFe]-Cofactor Biosynthesis in *Escherichia coli*"

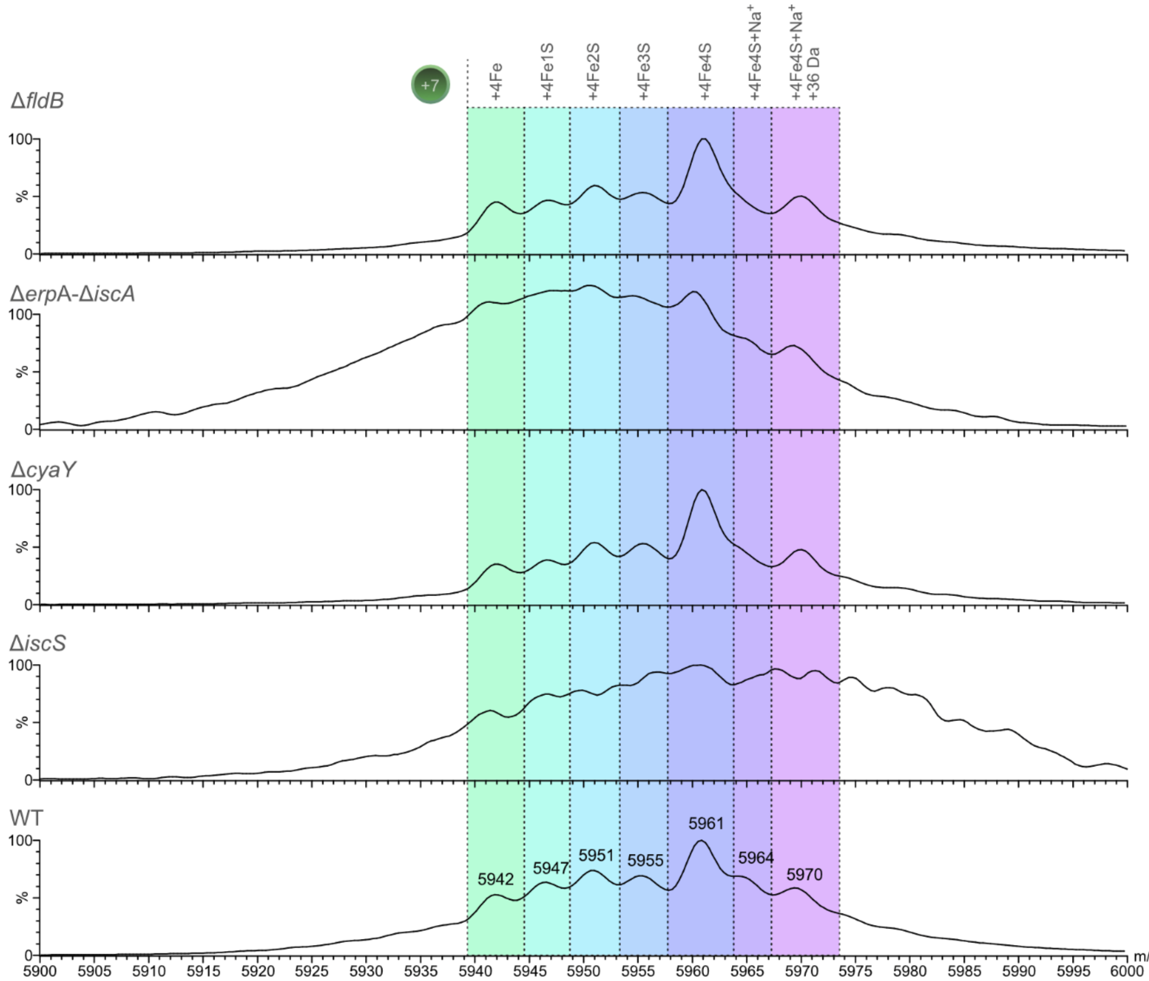


**Figure S1. Native mass spectra of [4Fe-4S] cluster-containing HypD dissociated from HypCD complexes isolated from different iron metabolism mutants.** Zoomed in native MS showing the +7 charged species of HypD including its [4Fe-4S] cluster dissociated from StrepII-tagged HypCD complex (charge state +12; collision energy 90V) isolated from strains MHD535 (Δ*cyaY*), PHB526 (Δ*iscS*), CP742 (Δ*iscA*-Δ*erpA*) and AS10 (Δ*fldB*) compared to the MC4100 (WT). All strains were transformed with plasmid pT-hypDCStrep.

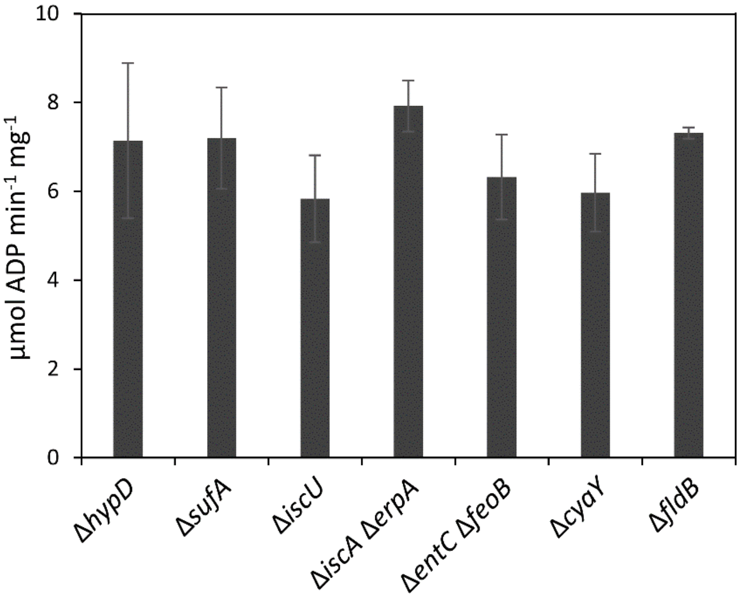


**Figure S2. ATPase activity of HypCD complexes is unaffected by *isc* mutations.** ATPase activity of purified, native HybG-HypD complexes isolated from the indicated mutants is shown. All strains carry plasmid pT-hypDCStrep encoding the StrepII-tagged HypCD complex.

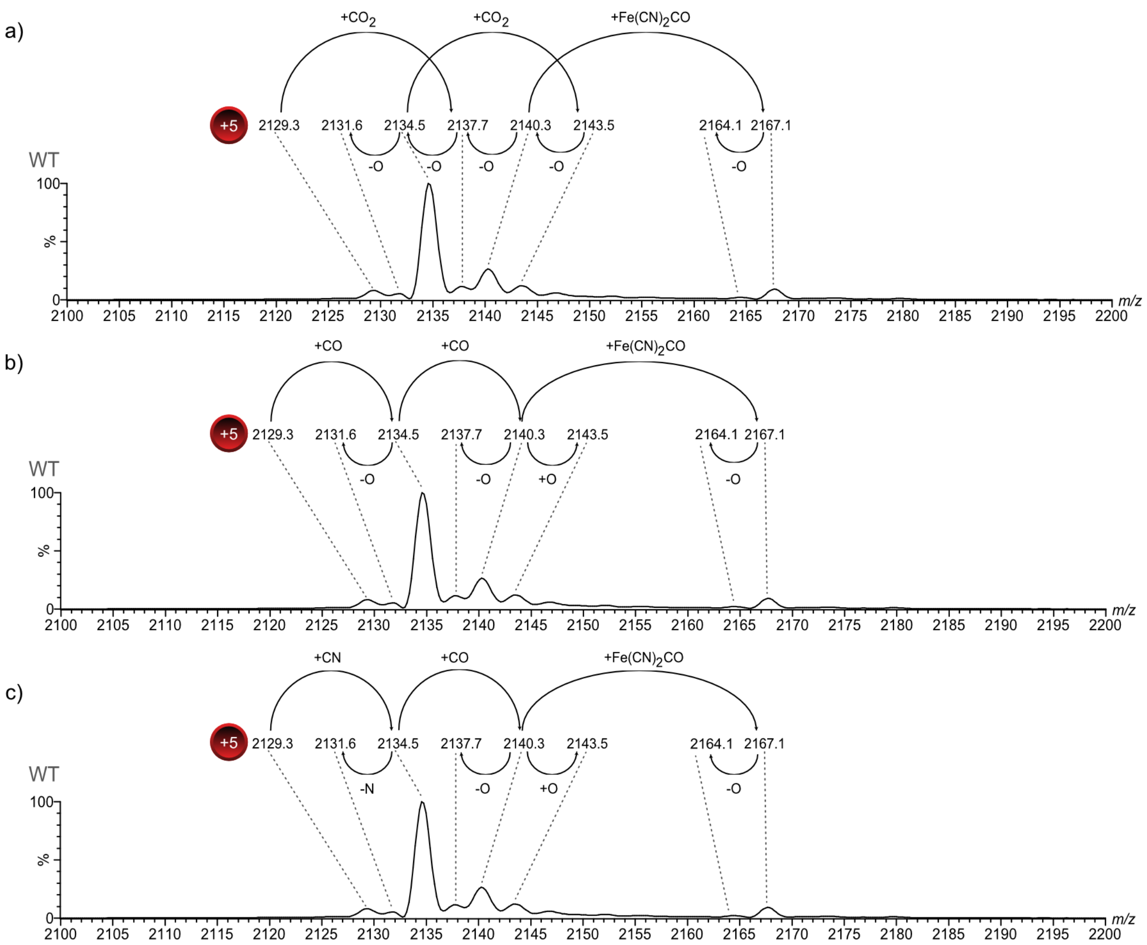


**Figure S3. Overview of potential adducts on HypC.** The three panels present different possible adducts accounting for the mass differences on HypC. – or +O signifies loss or addition of an oxygen. Note that the possibility of a methyl thiazolidine modification accounting for the +26 Da modification is not represented.

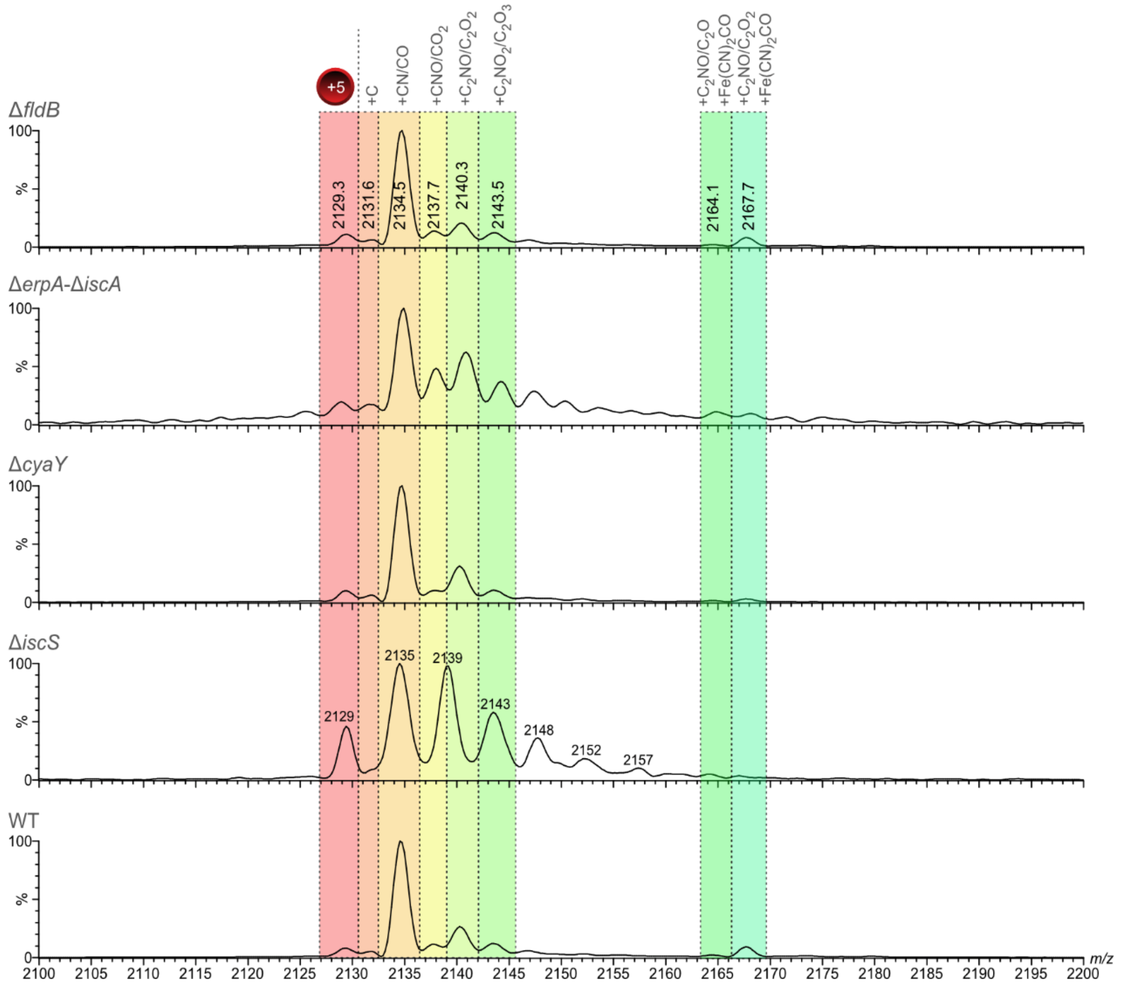


**Figure S4. Native mass spectra of HypC dissociated from HypCD complexes isolated from different iron metabolism mutants.** Zoomed in native MS showing the +5 charged species of HypC including its modifications dissociated from StrepII-tagged HypCD complex (charge state +12; collision energy 90V) isolated from strains MHD535 (Δ*cyaY*), PHB526 (Δ*iscS*), CP742 (Δ*iscA*-Δ*erpA*) and XYZ (Δ*fldB*) compared to the native HypCD complex (WT) isolated from DHP-D transformed with pT-hypDCStrep. Potential modifications are indicated above the colored overlay. Note that a putative methyl thiazolidine modification accounting for the +26 Da species is not indicated in the Figure.
